# Stress granules display bistable dynamics modulated by Cdk

**DOI:** 10.1101/2020.05.08.083782

**Authors:** G Yahya, AP Pérez, MB Mendoza, E Parisi, DF Moreno, C Gallego, M Aldea

## Abstract

Stress granules are conserved biomolecular condensates that originate in response to many stress conditions. These membraneless organelles contain nontranslating mRNAs and a diverse subproteome, but our knowledge on their regulation and functional relevance is still incipient. Here we describe a mutual-inhibition interplay between stress granules and Cdc28, the budding yeast Cdk. Amongst Cdc28 interactors acting as negative modulators of Start we have identified Whi8, an RNA-binding protein that localizes to SGs and recruits the mRNA of *CLN3*, the most upstream G1 cyclin, for efficient translation inhibition and Cdk inactivation under stress. However, Whi8 also contributes to recruiting Cdc28 to SGs, where it acts to promote their dissolution. As predicted by a mutual-inhibition framework, the SG constitutes a bistable system that is modulated by Cdk. Since mammalian cells display a homologous mechanism, we propose that the opposing functions of specific mRNA-binding proteins and Cdks subjugate SG dynamics to a conserved hysteretic switch.

## Introduction

Phase separation of proteins and nucleic acids into condensates is emerging as a general mechanism for cellular compartmentalization without the requirement of surrounding membranes (Banani *et al*, 2017; Alberti & Dormann, 2019; Shin & Brangwynne, 2017). These biomolecular condensates are based on weak multivalent interactions among component molecules that are mobile and exchange with the adjoining medium, and play essential roles in cell physiology as reaction crucibles, sequestration centers or organizational hubs. In a dynamic environment, cells need to control phase separation to form or dissolve condensates as a function of spatial and temporal cues. Thus, the molecular mechanisms that modulate phase separation will be critical to understanding how cells use biomolecular condensates to execute and control a growing list of cellular processes (Alberti *et al*, 2019; Snead & Gladfelter, 2019; Bratek-Skicki *et al*, 2020).

Stress granules (SGs) are conserved cytoplasmic condensates that contain (1) pools of nontranslating mRNAs and (2) a variety of proteins, including translation initiation factors and RNA-binding proteins that form core stable substructures in SGs (Protter & Parker, 2016). Non RNA-binding proteins such as post-translational modification factors, and protein or RNA remodeling complexes, are recruited to SGs by protein–protein interactions often mediated by intrinsically-disordered regions (IDRs), and modulate SG assembly and disassembly. However, a predominant role has been recently attributed to intermolecular RNA-RNA interactions as upstream determinants of stress granule composition (Van Treeck *et al*, 2018). Although SGs are thought to downregulate translation and protect recruited mRNAs in many different stress conditions, we still do not have sufficient experimental evidence to comprehend the relevance of SGs in cell physiology.

Stress restricts cell-cycle progression and budding yeast cells display a diverse set of mechanisms as a function of the stress signal. The HOG pathway constitutes a prominent paradigm, and operates on specific molecular targets to modulate different cell-cycle phases and transitions in response to osmotic stress (Solé *et al*, 2015; De Nadal *et al*, 2011). Regarding entry into the cell cycle, osmotic shock causes a temporary repression of the G1/S regulon (Bellí *et al*, 2001), in which Hog1-mediated phosphorylation of Whi5 and Msa1 contributes to inhibiting transcription (González-Novo *et al*, 2015). Downregulation of G1/S genes was also observed during heat and ER stress (Rowley *et al*, 1993; Vai *et al*, 1987) and, since chaperones play important but limiting roles at Start (Vergés *et al*, 2007; Yahya *et al*, 2014; Parisi *et al*, 2018), we proposed that, by compromising chaperone availability, all stressful conditions would target the chaperome as a common means to hinder entry into the cell cycle (Moreno *et al*, 2019). However, the precise molecular environment in which diverse stress conditions would converge to modulate Start is still unknown.

Here we describe the interplay between a common actor in stress, SGs, and the budding yeast Cdk, Cdc28. In a screen for Cdc28 interactors acting as negative modulators of Start we identified Whi8, a putative RNA-binding protein previously localized to SGs. Whi8 recruits the mRNA of *CLN3*, the most upstream G1 cyclin, to SGs and contributes to inhibiting its translation under stress conditions. On the other hand, Whi8-dependent recruitment of Cdc28 is important for timely dissolution of SGs when stress conditions are relieved. We also identified the key elements of a homologue mechanism in mammalian cells, and we propose that Cdks create a conserved bistable system that regulates SG dynamics.

## Results

### Whi8 is a Cdc28 interactor that hinders cell cycle entry

We previously identified and characterized a Cdc28^wee^ mutant that produces premature entry into the cell cycle causing a small cell-size phenotype and, to uncover new negative modulators of Start, we used quantitative proteomics to compare the interactomes of wild-type and wee Cdc28 proteins (Yahya *et al*, 2014) (Fig 1A). Among the group of proteins displaying a lower binding to Cdc28^wee^ we found YGR250c, a putative RNA-binding protein that was localized to stress granules (Buchan *et al*, 2008) and recently isolated as a multicopy suppressor of ER-mitochondria tethering complex defects (Kojima *et al*, 2016). To ascertain its role as a negative regulator of the yeast Cdk at Start we measured the budding volume of cells lacking YGR250c, and found a clear reduction that strictly required the presence of Cln3, the most upstream G1 cyclin (Fig 1B). Since overexpression of YGR250c produced the opposite effect and increased the budding volume by nearly 50%, we named YGR250c as Whi8, following the nomenclature of genes modulating cell size at Start. Similarly to Whi3 (Wang *et al*, 2004) and Whi7 (Yahya *et al*, 2014), loss or overexpression of Whi8 increased or decreased, respectively, the nuclear levels of Cln3 (Fig EV1).

**Figure 1.**
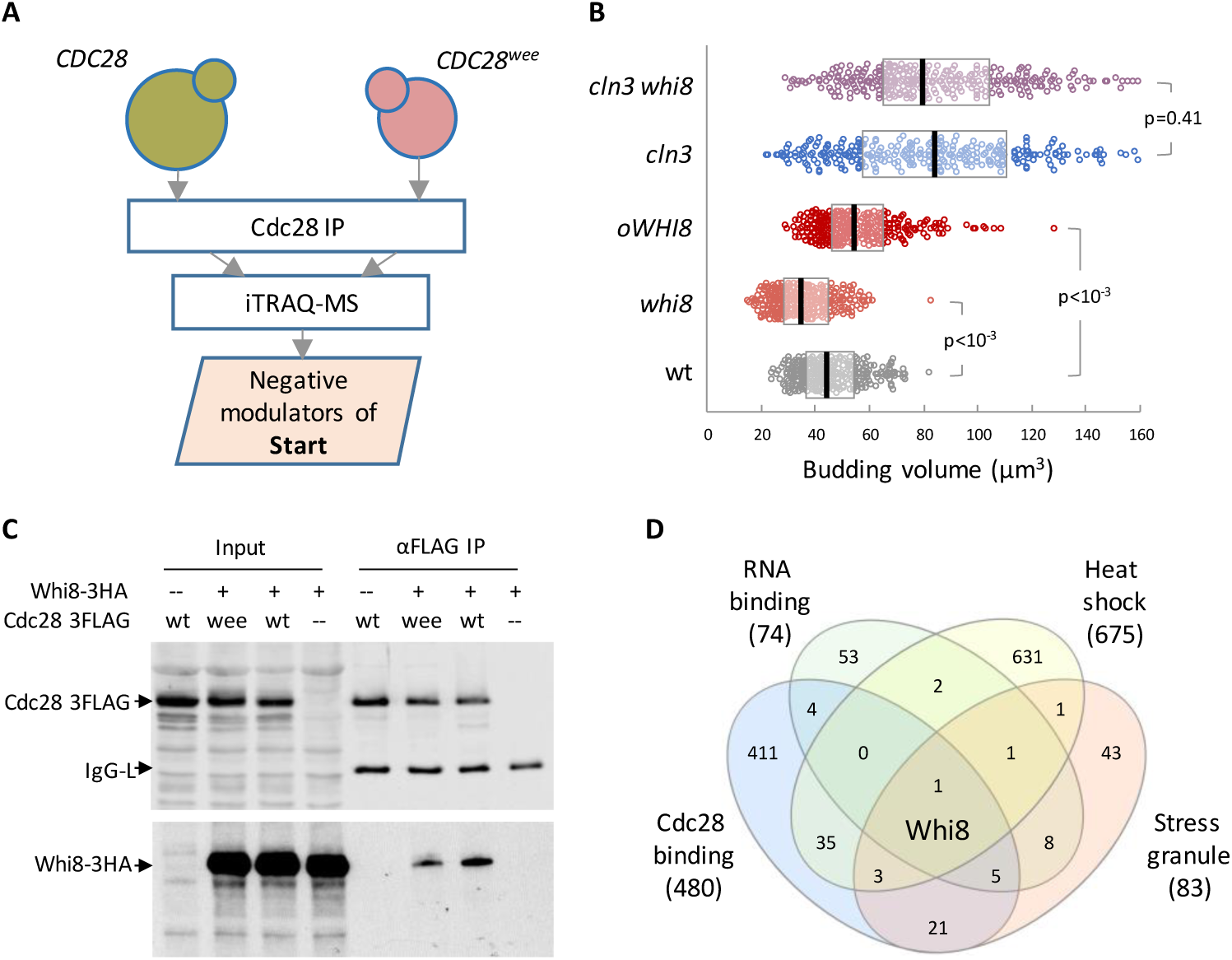
Whi8 is a Cdc28 interactor that hinders cell cycle entry. A Schematic of the screen for Cdc28 interactors as negative regulators of Start (Yahya *et al*, 2014). Briefly, pulldowns of wild-type and *wee* Cdc28 were analyzed by iTRAQ-based proteomics to identify proteins with a reduced affinity for Cdc28^*wee*^. B Cells with the indicated genotypes were analyzed to determine cell size at budding. Individual data (n>300) and median ± Q values are plotted. Shown p-values were obtained using a Mann-Whitney U test. C Immunoblot of input and αFLAG immunoprecipitation samples from *WHI8-3HA* cells expressing wild-type or mutant (*wee*) Cdc28-3FLAG proteins. D Venn diagram showing the overlapping of proteins that are (1) physical interactors of Cdc28, (2) contain demonstrated or putative RNA-binding domains, (3) are upregulated by heat shock and (4) were identified as SG components.

As expected, Whi8 levels were lower in Cdc28^wee^ immunoprecipitates compared to wild-type Cdc28 (Fig 1C), suggesting that the negative role of Whi8 at Start is mediated by physical interaction with the Cdc28 kinase. Notably, a bioinformatics analysis of 480 Cdc28 interacting proteins pinpointed Whi8 as the only gene product that (1) displays RNA-binding motifs, (2) is induced by heat shock and (3) is present in SGs (Fig 1D). In all, Whi8 emerged as a putative mediator of SG-born signals restraining entry into the cell cycle.

### Whi8 binds and recruits the *CLN3* mRNA to SGs

Since *cln3* was totally epistatic to the loss of Whi8 with regards to the cell size phenotype, we wanted to test whether the role of Whi8 at Start would be exerted on *CLN3*, perhaps due to its RNA-binding properties. We found that the *CLN3* mRNA was enriched 10-fold in Whi8 immunoprecipitates (Fig 2A), thus displaying a similar behavior to Whi3, a protein previously shown to bind the *CLN3* mRNA and regulate cell cycle entry (Garí *et al*, 2001; Colomina *et al*, 2008). Supporting shared roles in modulating *CLN3* expression by their RNA-binding motifs, Whi8 and Whi3 were found to co-immunoprecipitate in an RNA-dependent manner (Fig 2B). By contrast, the interaction between Whi8 and Pub1, a component of SGs, did not depend on RNA (Fig 2C).

**Figure 2.**
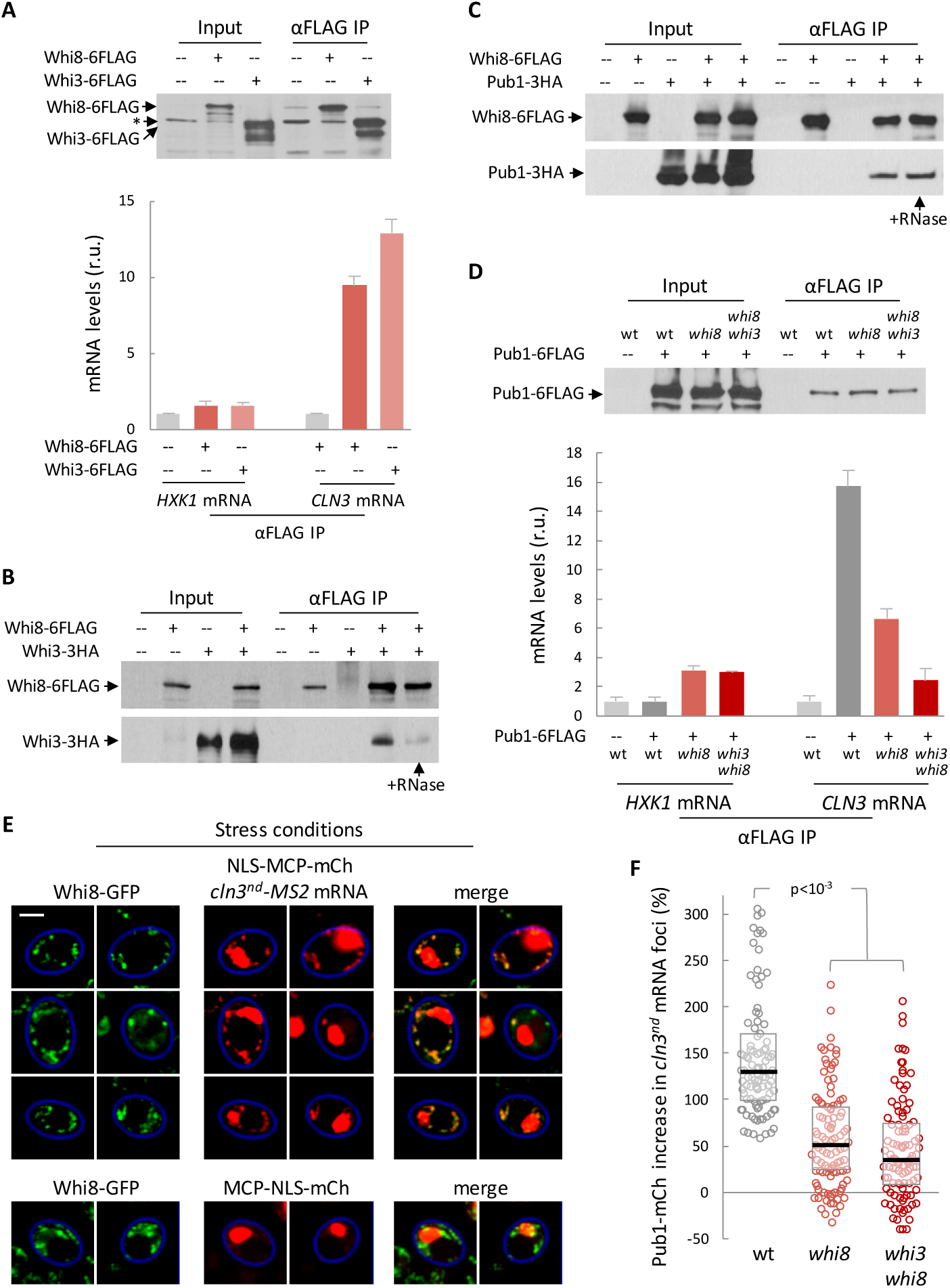
Whi8 binds and recruits the *CLN3* mRNA to SGs. A Immunoblot of input and αFLAG immunoprecipitation samples from *WHI8-6FLAG*, *WHI3-6FLAG* or untagged cells (top). Levels of *CLN3* and *HXK1* (as control) mRNAs in immunoprecipitates were determined and mean + SEM values (n=3) are plotted (bottom). B, C Immunoblots of input and αFLAG immunoprecipitation samples from cells expressing the indicated proteins. RNase was added during immunoprecipitation when indicated. D Immunoblot of input and αFLAG immunoprecipitation samples from wild-type, *whi8* or double *whi8 whi3* mutant cells expressing Pub1-6FLAG (top). Levels of *CLN3* and *HXK1* (as control) mRNAs were determined and mean + SEM values (n=3) are plotted (bottom). E Control (bottom) or *cln3^nd^-MS2* cells expressing Whi8-GFP and MCP-NLS-mCherry after 30 min at 42ºC in the absence of glucose. Bar, 2 µm. F Cells with the indicated genotypes expressing Pub1-mCherry, MCP-NLS-GFP and a *cln3*^*nd*^-*MS2* mRNA were stressed for 30 min at 42ºC in the absence of glucose, and the relative increase of Pub1-mCherry in *cln3*^*nd*^-*MS2* foci was measured. Individual data (n=100) and median ± Q values are plotted. Shown p-values were obtained using a Mann-Whitney U test.

The *CLN3* mRNA was clearly enriched in Pub1 pulldowns (Fig 2D) which, considering the abovementioned interactions, suggested that Pub1 could recruit the *CLN3* mRNA through Whi8. In agreement with this idea, enrichment of the *CLN3* mRNA in Pub1 pulldowns was strongly diminished in *whi8* cells and totally abrogated in double *whi8 whi3* cells (Fig 2D). We next tested whether the *CLN3* mRNA colocalized with Whi8 in SGs. To avoid the strong effects in cell size produced by high levels of Cln3, we expressed a *cln3*^*nd*^ mRNA that produces an inactive cyclin. As already described (Buchan *et al*, 2008), Whi8-GFP readily formed bright granules in the cytoplasm under stress conditions and, although the *cln3*^*nd*^ mRNA formed a much lower number of foci, most of them colocalized with Whi8-GFP granules (Fig 2E). Although the *CLN3* mRNA was previously found in SGs with Whi3 (Holmes *et al*, 2013; Cai & Futcher, 2013), the presence of Whi3 was totally dispensable. We found that colocalization of the *CLN3* mRNA with Pub1-mCherry granules required Whi8 and, to a much lesser extent, Whi3 (Fig 2F), which gives further support to the notion that the *CLN3* transcript is recruited to SGs with the essential role of Whi8.

### Whi8 is recruited to SGs via an IDR and is required to inhibit *CLN3* mRNA translation under stress conditions

Whi8 contains a C-terminal region of 120 residues largely dominated by disorder-prone amino acids (Fig 3A), suggesting that Whi8 could be recruited to SGs by an IDR. Deletion of these amino acids did not have any noticeable effect on cell size under normal growing conditions, but Whi8-ΔIDR was completely absent in the ER fractions as compared to the wild-type protein (Fig 3B). Moreover, although Whi8-ΔIDR was still recruited to SGs under stress conditions, the number of foci per cell and the fraction of Whi8-ΔIDR in SGs were sharply reduced (Fig 3C-F).

**Figure 3.**
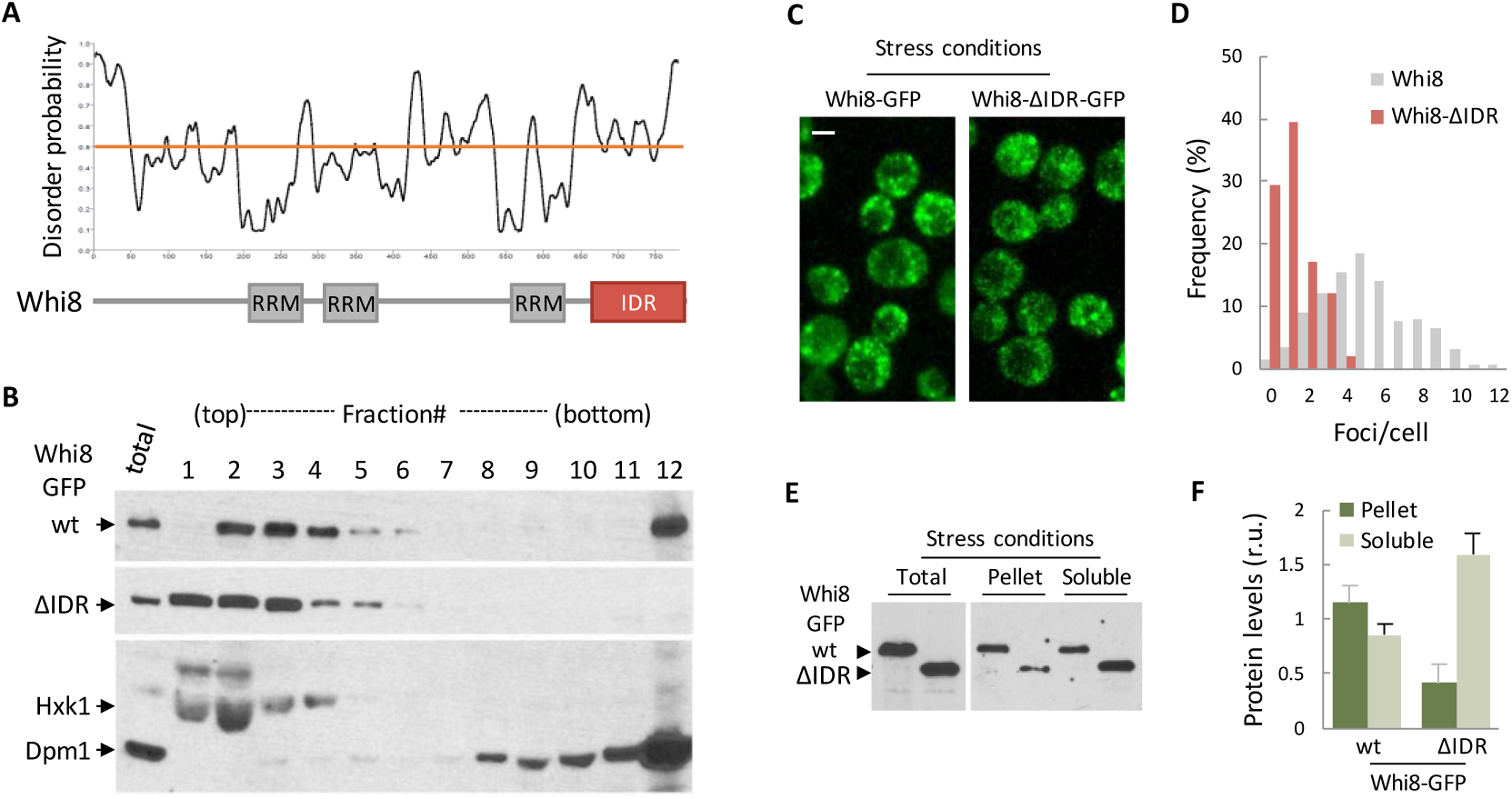
Whi8 contains an IDR important for SG recruitment. A Disorder probability of Whi8 amino acids and predicted RRMs and IDR. B Immunoblot analysis of the distribution of wild-type and ΔIDR Whi8-GFP in sucrose gradients. Hxk1 and Dpm1 are shown as soluble and ER markers, respectively. C Maximum projections of confocal images from cells expressing wild-type and ΔIDR Whi8-GFP after 30 min at 42ºC in the absence of glucose. Bar, 2 µm. D Cells expressing wild-type and ΔIDR Whi8-GFP were treated at 42ºC in the absence of glucose, and analyzed as in Fig 4A. Foci frequencies per cell are plotted. E Immunoblot of wild-type and ΔIDR Whi8-GFP in the indicated fractions from cells as in D. F Whi8-GFP levels as in **e** were quantified and mean + SEM values (n=3) are plotted.

One of the hallmarks of SG-recruited mRNAs is translation inhibition which, particularly for short-lived proteins such as Cln3, may have important effects in the corresponding downstream processes. We found that Cln3 levels rapidly decreased under stress conditions (Fig EV2A), reaching a five-fold reduction in less than 30 min, while the *CLN3* mRNA was only reduced by 40% compared to unstressed cells (Fig EV2C). Notably, levels of Cln3 decreased very slowly in cells lacking Whi8 (Fig EV2A), thus reinforcing the notion that Whi8 would recruit the *CLN3* mRNA to SGs to inhibit its translation under stress conditions. Overexpression of SG components promotes SG formation and recapitulates inhibition of SG-reporter gene expression in non-stressed cells (Kedersha & Anderson, 2007). Accordingly, when we overexpressed Whi8 in the absence of stress we observed a significant decrease in Cln3 protein levels (Fig EV2B), while mRNA levels remained unchanged (Fig EV2C). These data would explain the observed large cell-size phenotype and low Cln3 levels in the nucleus of Whi8-overexpressing cells, and emphasize the idea that Whi8 acts as a negative regulator of Start by modulating expression of the *CLN3* mRNA.

### Cdc28 is recruited to SGs and modulates SG dynamics

Cdc28 has been colocalized with Pbp1 in foci of stationary-phase cells (Shah *et al*, 2014) and, although not quantitative, a proteomic survey reported the presence of Cdc28 in SGs (Jain *et al*, 2016). To ascertain this possibility, we analyzed Cdc28-GFP and Pub1-mCherry in live cells under stress conditions and found that both proteins displayed extensive colocalization patterns (Fig 4A). Careful measurement of the relative levels of the two proteins in SGs yielded a steeper slope when Cdc28-GFP was taken as a dependent variable (Fig 4B), favoring Pub1 as an upstream factor for Cdc28 localization to SGs. Notably, the ratio of Cdc28-GFP to Pub1-mCherry in SGs decreased to ca. 40% when comparing *whi8* with wild-type cells (Fig 4C). Similarly to other components of the SG (Youn *et al*, 2018), Cdc28 and Pub1 were found to co-immunoprecipitate under non-stress conditions (Fig 4D). In all, these data underscore the relevance of Whi8 and Pub1 in recruiting Cdc28 to SGs.

**Figure 4.**
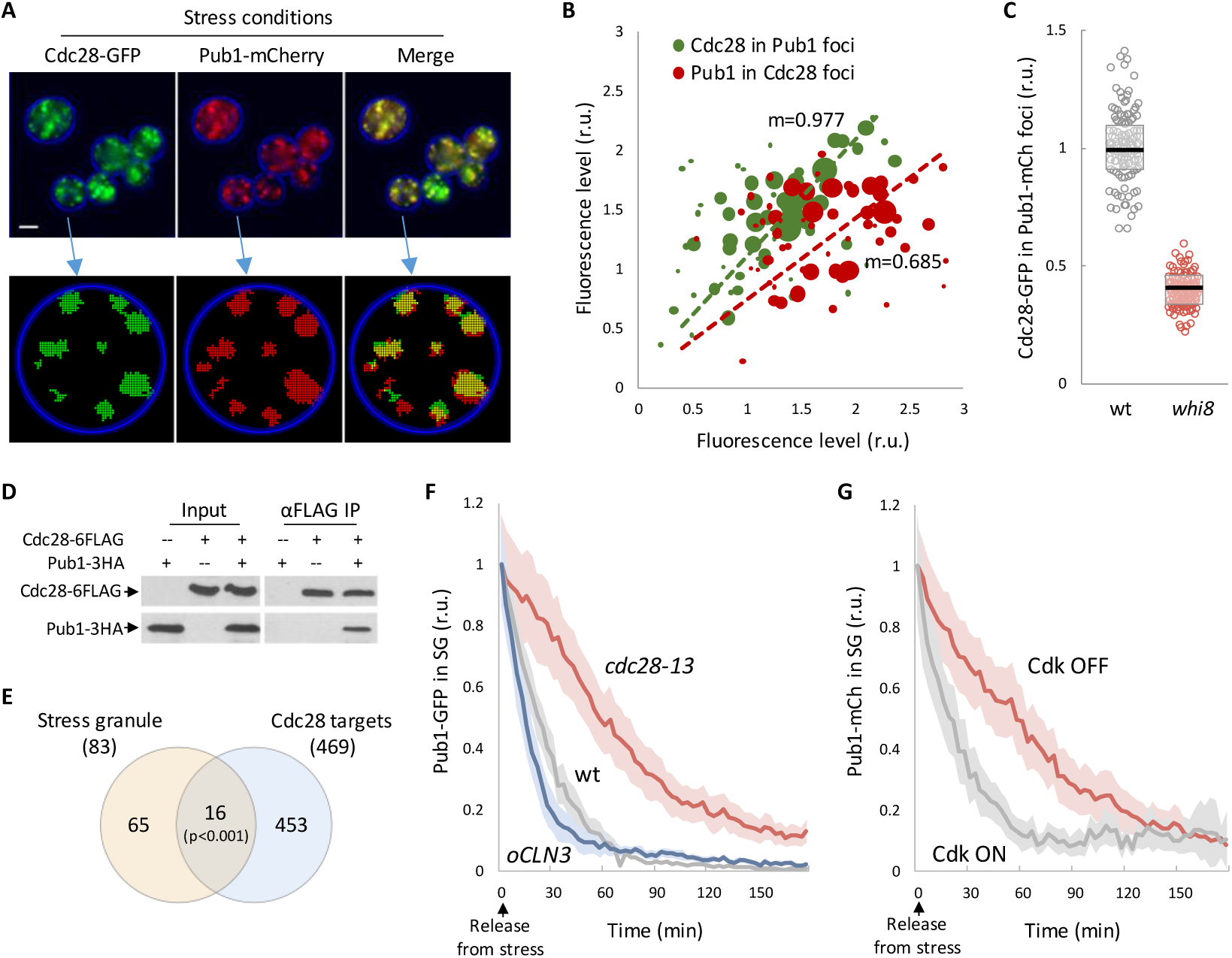
Cdc28 is recruited to SGs and modulates SG dynamics. A Maximum projections of confocal images from cells expressing Cdc8-GFP and Pub1-mCherry after 30 min at 42ºC in the absence of glucose (top). Bar, 2 µm. GFP and mCherry foci detected for quantification in a sample cell with the aid of BudJ are also shown (bottom). B Cdc8-GFP levels in Pub1-mCherry foci (n=60), and vice versa, from cells as in (A) are plotted with point diameter corresponding to foci size. Slope (m) values are also shown. C Cdc8-GFP levels in Pub1-mCherry foci were measured in wild-type and *whi8* cells stressed for 30 min at 42ºC in the absence of glucose. Individual data (n=125) and median ± Q values are plotted. D Immunoblot of input and αFLAG immunoprecipitation samples from cells expressing Cdc28-6FLAG and Pub1-3HA. E Venn diagram showing the overlapping of proteins that (1) were identified as SG components and (2) are putative phosphorylation targets of Cdc28. The indicated p-value was obtained assuming completely independent allocations. F Wild-type, *cdc28-13* and *oCLN3* (*tetO_2_-CLN3*) cells expressing Pub1-mCherry were stressed for 30 min at 42ºC in the absence of glucose and, once released at 37ºC in the presence of glucose, Pub1-mCherry levels in foci were measured at different time points. Mean values (n>30) and confidence limits for the mean (α=0.05) are plotted. G *GALp-CLN3 cln1,2* expressing Pub1-mCherry were grown in galactose and arrested in G1 by transfer to glucose for 2 hr (Cdk OFF). Cells were then stressed for 30 min at 42ºC in the absence of glucose and, once released at 30ºC in the presence of glucose to maintain the G1 arrest, Pub1-mCherry levels in foci were measured at different time points. As control, *CLN3 cln1,2* cells (Cdk ON) were subject to the same experimental conditions. Mean values (n>30) and confidence limits for the mean (α=0.05) are plotted.

A significant number of SG proteins (Jain *et al*, 2016) may be phosphorylation targets of Cdc28 (Fig 4E) and, in addition to Whi8, we found several translation initiation factors that are thought to be involved in translational repression in SGs (Fig EV3A). This observation led us to hypothesize that Cdc28 could play a role in SG assembly and dissolution. To test this possibility, we carefully measured the dissolution kinetics of Pub1-mCherry foci by time-lapse microscopy, and first compared cells carrying wild-type *CDC28* and thermosensitive *cdc28-13* alleles. Glucose was added to induce release from stress, but the temperature was only decreased to 37ºC to maintain a restrictive scenario for the *cdc28-13* allele. As shown in Fig 4F, wild-type cells readily dissolved SGs under these partial-release conditions, and the levels of Pub1-mCherry in SGs decreased to 50% in only 21 min. By contrast, SG dissolution was much slower in *cdc28-13* cells, taking longer than 60 min to reach a 50% reduction in Pub1-mCherry in SGs (Fig 4F). As an independent approach we used a G1-cyclin conditional strain that lacks Cln1,2 and holds Cln3 under the control of a regulatable promoter, which causes cells to arrest in G1 with no Cdk activity under repression conditions. We found a strikingly similar delay in SG dissolution when comparing G1-arrested and cycling cells (Fig 4G), which confirms the key role of Cdk activity in SG dissolution. Giving further support to this notion, cycling wild-type cells displayed clear differences in SG dissolution kinetics depending on cell cycle position, being slower when Cdk activity is lower, i.e. G1 phase (Fig EV3B). Finally, overexpression of Cln3 produced the opposite effects and reduced by 30% the half-life of SGs after release form stress (Fig 4F). Albeit surprisingly, the presence of *CLN3-1*, a truncated hyperstable allele that reproduces most phenotypes of *CLN3* overexpression, was much less effective in SG dissolution (Fig EV3C). Cln3-1 lacks a putative 91-aa IDR that could play additional roles in recruiting the Cdc28/Cln3 complex to SGs.

### Mammalian SGs contain Cdk-cyclin factors and are modulated by cell cycle position

SGs from yeast and mammalian cells display a highly significant overlap in composition and share distinct substructural traits (Jain *et al*, 2016). Thus, we anticipated that our findings in yeast cells would also apply to mammalian SGs. To test whether G1 cyclin mRNAs are recruited to mammalian SGs we used the same MS2-based approach as in yeast and observed a clear co-localization of a *CCND1-MS2* mRNA with TIA1, the Pub1 mammalian homologue, in SGs of HeLa cells (Fig EV4A). It was shown that Caprin1, a component of SGs, binds the cyclin D2 mRNA in non-stressed 293T cells (Solomon *et al*, 2007). We found that Caprin1 and *CCND1-MS2* mRNA also colocalized in SGs (Fig 5A). Interestingly, shRNA-driven downregulation of Caprin1 levels was accompanied by a reduction in the number of foci produced by the *CCND1-MS2* mRNA under stress conditions (Fig 5B), suggesting that Caprin1 could act as a functional homologue of Whi8 in mammalian cells, where cyclin D1 protein levels are also strongly downregulated during SGs formation (Fig EV4B).

**Figure 5.**
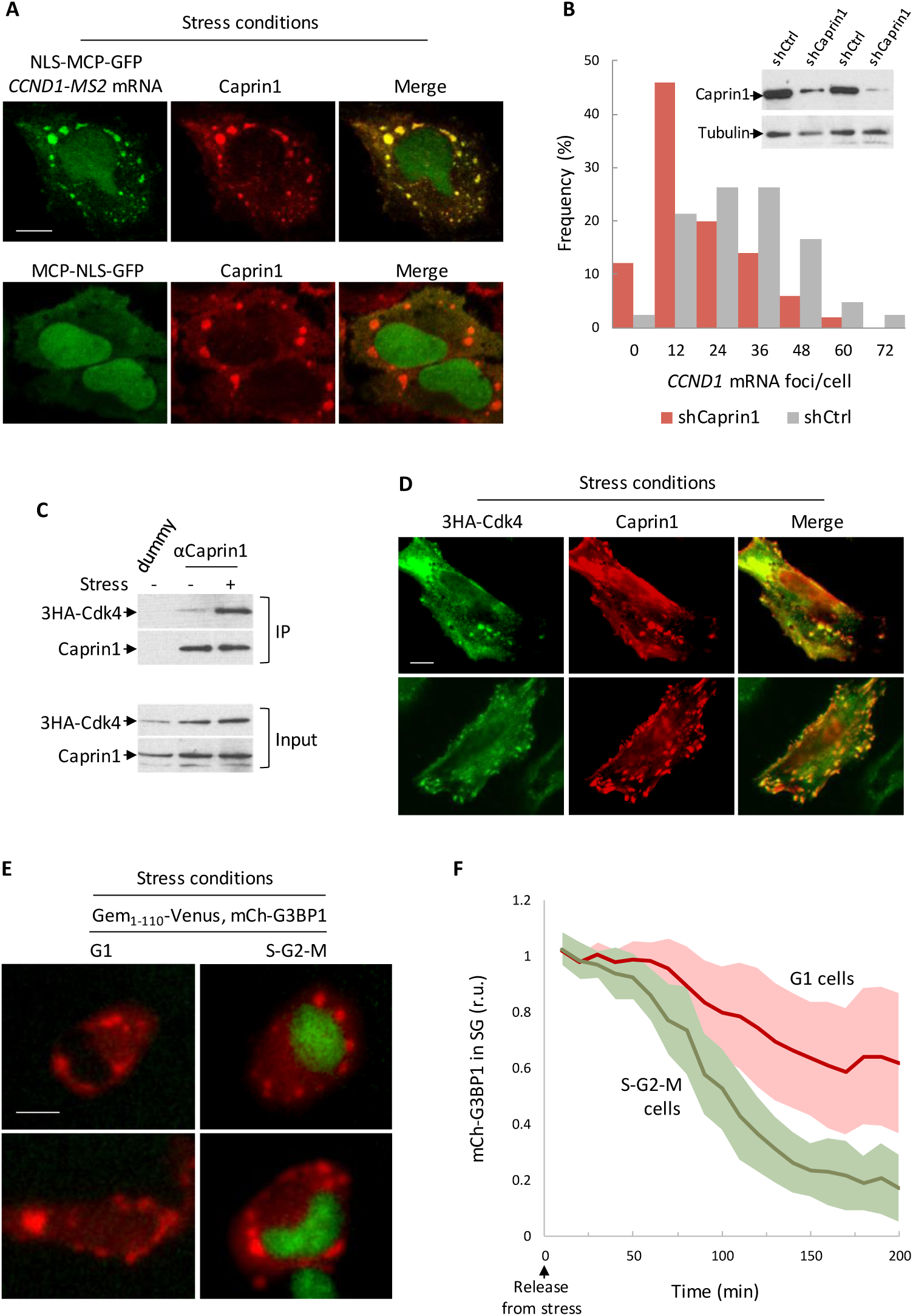
Mammalian SGs contain Cdk-cyclin factors and are modulated by cell cycle position. A HeLa cells expressing NLS-MCP-GFP and either a *CCND1-MS2* mRNA or none (as control) were subject to 0.5 mM NaAsO_2_ for 30 min and analyzed by immunofluorescence with a αCaprin1 antibody. Bar, 5 µm. B HeLa cells expressing NLS-MCP-GFP, a *CCND1-MS2* mRNA and either shCaprin1 or shCtrl were subject to 0.5 mM NaAsO_2_ for 30 min, and analyzed as in Fig 4A. Foci frequencies per cell are plotted. Inset: Immunoblot analysis of Caprin1 levels in total extracts of HeLa cells expressing shCaprin1 or shCtrl. Tubulin serves as loading control. C Immunoblot of input and αCaprin1 immunoprecipitation samples from HeLa cells expressing 3HA-Cdk4 in the presence (+) or absence (-) of 0.5 mM NaAsO_2_ for 30 min. D HeLa cells expressing 3HA-Cdk4 were subject to 0.5 mM NaAsO_2_ for 30 min and analyzed by immunofluorescence with αHA and αCaprin1 antibodies. Bar, 5 µm. E Representative images of U2OS cells expressing Geminin_1-110^−^_mVenus and mCherry-G3BP1 after treatment with 0.5 mM NaAsO_2_ for 30 min. Bar, 5 µm. F U2OS cells expressing Geminin_1-110_-mVenus and mCherry-G3BP1 were treated with 0.5 mM NaAsO_2_ for 30 min as in (E) and, once released in fresh medium, mCherry-G3BP1 levels in foci were measured at different time points. Mean values (n=25) and confidence limits for the mean (α=0.05) are plotted.

Next, we decided to test whether Caprin1 and Cdk4, a G1 Cdk, interact by co-immunoprecipitation as in yeast cells, and found that these proteins are bound in a stress-dependent manner in HeLa cells (Fig 5C). Moreover, we observed almost overlapping patterns of localization of Caprin1 and Cdk4 in SGs (Fig 5D).

The rate of SG dissolution has been shown to be strongly diminished by Cdk inhibitors in HeLa and U2OS cells (Wippich *et al*, 2013). Hence, the presence of a G1 Cdk and the Caprin1-mediated recruitment of a G1 cyclin mRNA in SGs suggested that cell cycle progression could also play a role in SG dynamics in mammalian cells. As in yeast cells, Cdk activity is low in G1, suddenly increases during the G1/S transition, and becomes maximal during mitosis until anaphase. Using an mCherry-G3BP1 reporter to analyze SG dissolution kinetics in U2OS cells expressing a mVenus-Gem^1-110^ fusion that is stable from S phase to anaphase, we found that SG half-life was reduced by 2-fold in cells progressing through S-G2-M phases, when Cdk activity is high, compared to G1 cells (Fig 5E,F). These data confirm our findings in yeast cells and point to the notion that SGs would antagonize Cdk activity by restraining cyclin translation under stress conditions and that, in turn, Cdk would accelerate SGs dissolution during release from stress.

### Mutual inhibition as a bistable system for SG dynamics

To gain insight into the relevance of the counteracting effects between SGs and Cdk activity, we modeled SG dynamics in a mutual-inhibition system with Cdk (Fig 6A and Fig EV5A). Briefly, we assumed that active Cdk (*Cdk*_*Cyc*_) would act as an enzyme with specific catalytic (*k*_*c*_) and Michaelis-Menten (*K*_*M*_) constants to promote dissolution of SG factors (*SG*) as free components (*SG*_*Comp*_). All other processes were defined by simple mass-action laws with explicit rate constants. We tried to keep the model as simple as possible in order to obtain all the kinetic parameters from experimental data. The *K*_*M*_ for Cdc28 was previously determined *in vitro* (Bouchoux & Uhlmann, 2011), and the maximal substrate concentration in SGs was estimated as described in Fig EV5B. We obtained the basal SG-dissolution rate constant (*k*_*bd*_) by fitting the model to SG dissolution in the *cdc28-13* mutant in the absence of stress (Fig EV5C), where all other variables have no effect. We then used SG formation and dissolution data from wild-type cells to estimate the remaining rate constants (Fig EV5D). With these experimental parameters, the mutual-inhibition model predicted a bistable switch as a function of the stress signal (Fig 6B). To observe how the bistability of the model arises, in Fig 6C we plotted the steady-state levels of condensed SG factors (*SG*) and active Cdk (*Cdk*_*Cyc*_) as independently predicted by each of the two modules of the system, either (1) *SG* dissolution as a function of *Cdk*_*Cyc*_ and the stress level (*Stress*) or (2) *Cdk*_*Cyc*_ inactivation as a function of *SG*.

**Figure 6.**
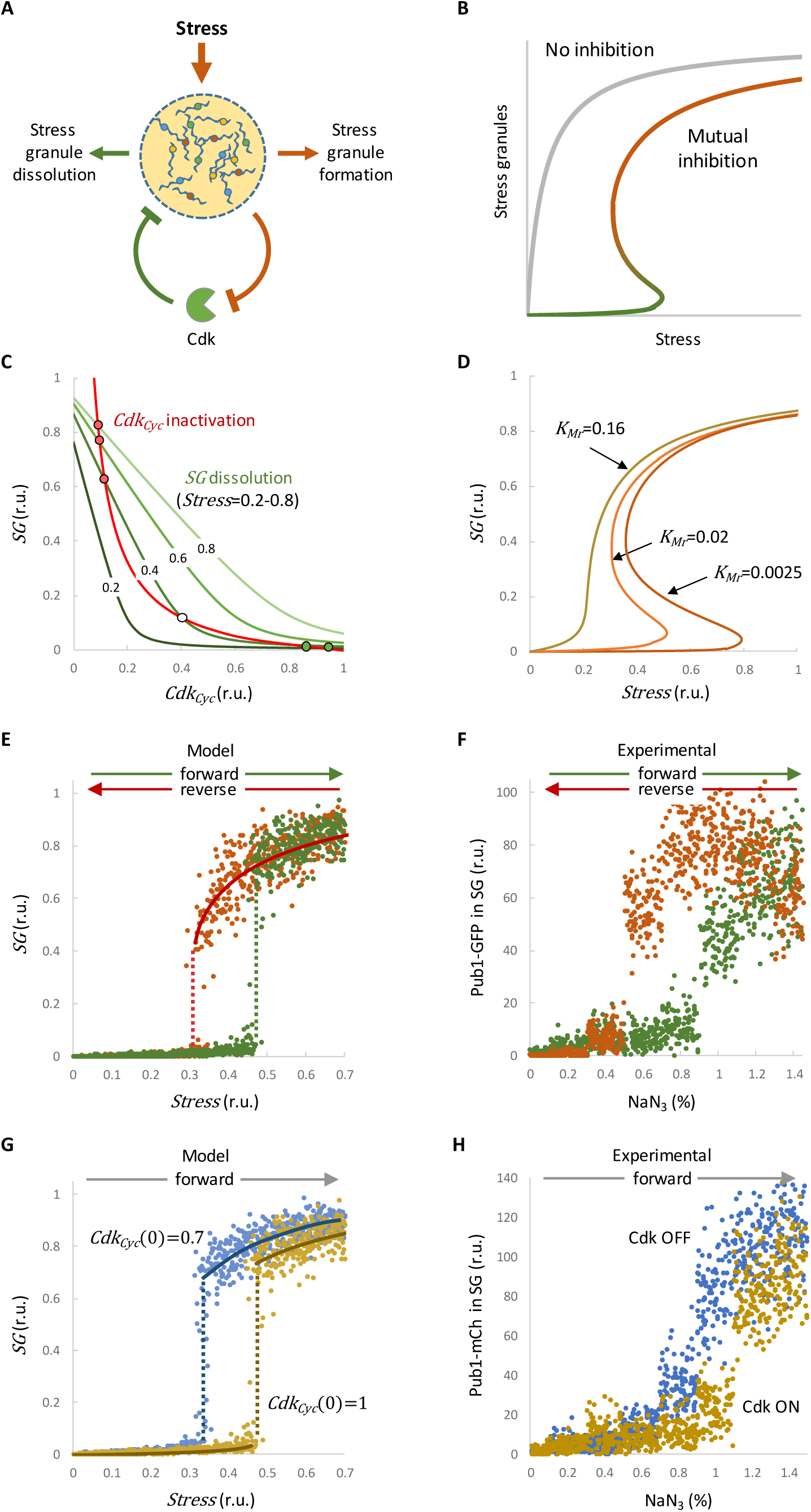
Mutual inhibition as a bistable system for SG dynamics. A SGs maintain cyclin mRNAs translationally inactive and, hence, inactivate the Cdk. In turn, active Cdk promotes SG dissolution and cyclin mRNA release and translation, thus accelerating SG disassembly. B SG and Cdk mutual inhibition creates a bistable system that elicits SG formation only above a certain degree of stress and maintains SG integrity until normal conditions are almost fully restored. C Steady-state balance plots of *SG* and *Cdk*_*Cyc*_ as derived from *Stress*-modulated SG dissolution by active Cdk (green lines) or Cdk inactivation by SGs (red line) reactions. Unstable (white circle) and stable (red and green circles) steady states are indicated. D *SG* steady states vs *Stress* as a function of the *K*_*Mr*_. E Simulations of *SG* levels in forward or reverse modes. The plot shows final *SG* steady states when the initial *Stress* variable was set to 0 (forward mode, green) or set to 1 (reverse mode, red), simulating non-stressed and stressed cells, respectively. Both deterministic (lines) and stochastic (dots) simulations are shown. F SG steady-state levels in forward or reverse mode experiments. In forward mode (green), exponentially-growing wild-type cells expressing Pub1-GFP were subject to different NaN_3_ concentrations for 60 min. In reverse mode, cells were first subject to 1.4% NaN_3_ for 60 min, and then incubated in the presence of different NaN_3_ concentrations for an additional 60-min period. Pub1-GFP levels in foci were measured in single cells (n>300) and bootstrapped (n=50) mean values are plotted. G Simulations of *SG* levels as a function of the total Cdk levels in forward mode. The plot shows final *SG* steady states when the initial *Cdk*_*Cyc*_ variable was set to 1 (yellow) or reduced to 0.7 (blue). Both deterministic (lines) and stochastic (dots) simulations are shown. H SG steady-state levels in forward mode experiments with cycling and G1-arrested cells with no Cdk activity. *GALp-CLN3 cln1,2* expressing Pub1-mCherry were grown in galactose and arrested in G1 by transfer to glucose for 2 hr (Cdk OFF). Then, cells were stressed for 60 min in the presence of different NaN_3_ concentrations. As control, *CLN3 cln1,2* cells (Cdk ON) were subject to the same experimental conditions. Pub1-mCherry levels in foci were measured in single cells (n>300) and bootstrapped (n=50) mean values are plotted.

At low stress, *SG* dissolution and *Cdk*_*Cyc*_ inactivation curves intersect only once and the system is monostable, with high active Cdk1 and low condensed SG factors. When stress reaches a certain value, the system creates a saddle-node bifurcation and a new stable steady state, with low active Cdk1 and high levels of condensed SG factors. At higher stress levels, the *SG* dissolution curve is pushed further and the system becomes monostable again, with low active Cdk1 and high SG condensation (Fig 6C). The relative *K*_*M*_, which incorporates SG component concentration, has strong effects on the predicted bistability (Fig 6D), thus highlighting the importance of substrate accumulation in the SG itself to attain a bistable system.

We then tested whether SG steady states followed a pattern compatible with bistability as a function of stress. As a tunable stress effector we used NaN_3_ at different concentrations and measured SG formation after 1h of treatment. We found that steady-state SG levels increased with stressor concentration following bistable kinetics, and displayed a hysteretic behavior as predicted by the model when stressor concentration was reversed (Fig 6E,F). Finally, we used the G1-cyclin conditional strain to test the effects of Cdk activity in the switch-like behavior of SG formation as a function of NaN_3_ concentration. Although a sigmoidal curve was still observed, G1-arrested cells with no Cdk activity advanced SG formation at lower stress levels in a similar fashion to what the model predicts if kinase levels are decreased (Fig 6G,H). These data support the important role of Cdk in SG dynamics, and suggest that other kinases or factors important for SG formation and dissolution also act in mutual-inhibition modules.

## Discussion

Here we identified Whi8 (YGR250C) as a protein that interacts with the Cdc28 Cdk and recruits the *CLN3* cyclin mRNA to SGs for translational repression and, hence, Cdc28 inactivation under stress conditions in G1 cells. Moreover, the yeast Cdk is also recruited to SGs with the important participation of Whi8, and plays a crucial role in SG dissolution when cells are returned to non-stress conditions. We found a similar scenario in mammalian cells, where the *CCND1* cyclin mRNA is translationally repressed by stress and recruited to SGs with the contribution of Caprin1, an RNA-binding protein that interacts with Cdk4, a G1 Cdk, in a stress-dependent manner. While Cdk4 colocalizes with Caprin1 in SGs, SG dissolution is slower in cells where Cdk activity is lower, i.e. G1 cells, when released from stress conditions. Thus, Whi8 and Caprin1 would act as molecular links between Cdk inactivation and Cdk-dependent SG dissolution during adaptation to and recovery from stress, respectively. Finally, our data show that SGs behave as a Cdk-dependent bistable system that only switches when stress levels reach a minimal threshold or normal conditions are almost completely restored.

Recent work analyzed the presence of mRNAs in yeast SGs (Khong *et al*, 2017), and found that G1 cyclin mRNAs were enriched in the SG fraction, 4-fold for *CLN3* and about 2-fold for *CLN1,2*, but levels of G1 cyclin mRNAs were still significant in the soluble fraction. Moreover, similar data from mammalian cells showed no or only a modest enrichment of *CCND1,2,3* mRNAs in SGs (Khong *et al*, 2017; Namkoong *et al*, 2018). Thus, although recruitment to SGs likely plays an important role, other mechanisms such as eIF2α inhibition by phosphorylation (Crawford & Pavitt, 2019) could act to ensure full inhibition of translation of G1 cyclin mRNAs under stress conditions. On the other hand, recruitment of a fraction of cyclin mRNAs to SGs could increase local Cdk activation when translation resumes after release from stress.

A very significant number of proteins belonging to SGs may be phosphorylated by Cdc28 in yeast cells and, not surprisingly, most of them have functional homologues in mammalian cells (Fig EV3A). The reason Cdc28 would have such a high number of targets in SGs likely resides in their dependence on multivalent protein-protein interactions, which also agrees with the fact that no single protein has been found to be essential for SG formation (Yoon *et al*, 2010; Yang *et al*, 2014; Buchan *et al*, 2013). Intriguingly, a comparative analysis of Cdc28-target phosphosites revealed that their position, rather than being conserved, is very dynamic within disordered regions (Holt *et al*, 2009).

We show that SG dynamics obey a bistable system where the Cdk is an important effector, and we have recapitulated this behavior with a simple mutual-inhibition model. Notably, the K_M_ to substrate ratio was crucial to attaining bistability and, if Cdk substrates were always in a soluble form in the cytoplasm, the model would only predict monostable steady-states at varying stress levels. In other words, the higher substrate concentration attained in SGs is what decreases the K_M_/substrate ratio and makes the model bistable. Thus, the effects of mutual-inhibition would be especially relevant during SG dissolution, rather than SG assembly. If this were the case, SG components would not be necessarily phosphorylated when soluble in the cytoplasm under normal conditions and, hence, the Cdc28 phosphoproteome could include many other SG proteins in addition to those listed in Fig EV3A. Alternatively, since many SG components display physical interactions even in the absence of stress (Youn *et al*, 2018), they could interact with Cdc28 in supramolecular complexes to increase their effective concentration as substrates also under non-stress conditions. Favoring the later possibility, we found that Cdc28 and Whi8 co-immunoprecipitate vey efficiently in the absence of stress.

SG dissolution requires the Cdc48 segregase in yeast (Buchan *et al*, 2013) and mammalian (Wang *et al*, 2019) cells, and we previously found that Cdc28 phosphorylates Cdc48 to enhance its segregase activity in releasing Cln3 from the ER during G1 (Parisi *et al*, 2018). Therefore, Cdc28 could also act on SG dissolution by modulating the affinity and/or segregase activity of Cdc48 towards components of the SG.

Giving support to our results, a screen in mammalian cells had pinpointed Cdk inhibitors by their marked effects in delaying SG dissolution (Wippich *et al*, 2013). These authors identified Dyrk3 as a key factor modulating SG dynamics, and recently showed that this kinase is also important for dissolution of specific membraneless organelles during mitosis (Rai *et al*, 2018). Also in mammalian cells, SG dissolution is modulated by FAK-dependent phosphorylation of Grb7 (Tsai *et al*, 2008). In yeast, Sky1 is recruited to SGs, where it phosphorylates Npl3 and modulates their dynamics (Shattuck *et al*, 2019). Therefore, fast and efficient SG assembly and dissolution would result from the concerted action of different protein kinases. Nonetheless, the unique mutual inhibitory roles of Cdk and SGs would provide bistability and hysteresis to prevent SG formation at low levels of stress and sustain their presence until normal conditions are entirely restored. Indeed, as our data suggest, the SG-Cdk mutual inhibitory scenario should also apply to other condensation modulators, thus subjugating SG dynamics to a robust switch as a function of different facets of the cellular physiological status.

## Acknowledgements

We thank E. Rebollo for technical assistance, and B. Futcher, R. Singer and T. Stracker for providing strains, cell lines and plasmids. We also thank C. Rose for editing the manuscript. This work was funded by the Spanish Ministry of Science and Innovation, and the European Union (FEDER) to C.G. (BFU2017-83375-R) and M.A. (BFU2016-80234-R). A.P.P. and D.F.M. received FI fellowships of *Generalitat de Catalunya*.

## Author contributions

G.Y., A.P.P., M.M.B., E.P. and D.F.M. built genetic constructs and strains, and performed the experiments. M.A. implemented the mathematical model and performed the informatics analysis. C.G. and M.A. conceived the study, analyzed the data and wrote the manuscript.

## Materials and Methods

### Cells and growth conditions

Yeast strains and plasmids are listed in Tables EV1 and EV2, and methods used for chromosomal gene transplacement and PCR-based directed mutagenesis were previously described (Ferrezuelo *et al*, 2012). With the exception of overexpression experiments, all gene fusions in this study were expressed at endogenous levels. Cells were grown for 7-8 generations in SC medium with 2% glucose at 30ºC unless stated otherwise. *GAL1*p-driven gene expression was induced by addition of 2% galactose to cultures grown in 2% raffinose at OD600=0.5 or, in the presence of glucose, by adding 1 µM β-estradiol to cells expressing the Gal4-hER-VP16 transactivator (Louvion *et al*, 1993). When indicated, overexpression of *CLN3* was performed with a *tetO*-based expression system (Garí *et al*, 1997). SG formation was routinely induced by transferring cells to 42ºC for 30 min in SC medium without glucose, and SG dissolution was assessed by returning stressed cells to 30ºC in SC medium supplemented with glucose. In experiments shown in Fig 4F, SG dissolution was assessed at 37ºc to maintain restrictive conditions for the thermosensitive *cdc28-13* allele. Finally, SG steady states were analyzed by treating cells with the indicated concentrations of NaN_3_ for 60 min at 30ºC in SC plus 1% glucose.

Human HeLa and U2OS cells were grown at 37ºC in Dulbecco’s modified Eagle’s medium (DMEM) supplemented with glutamine, antibiotics and 10% FBS. Transfection with Caprin1-directed or control shRNAs and vectors expressing NLS-MCP-GFP, 3HA-Cdk4, Geminin_1-110-_mVenus or mCherry-G3BP1 proteins, or *CCND1-MS2* mRNA, was performed as described (Ruiz-Miro *et al*, 2011). Cycling cells were analyzed 18 hr after replating, and SG formation was induced by addition of 0.5 mM NaAsO_2_ for 30 min, unless stated otherwise. SG steady states were analyzed after 60 min in NaAsO_2_ at the indicated concentrations.

### Subcellular fractionation, immunoprecipitation and immunoblot analysis

Subcellular fractionation was as described (Vergés *et al*, 2007). FLAG-tagged proteins were immunoprecipitated with αFLAG (clone M2, Sigma) beads (Wang *et al*, 2004), and immunoblot analysis was performed as described (Gallego *et al*, 1997). Used antibodies are listed in Table EV3.

### Time-lapse wide-field and confocal microscopy

Cells were analyzed by time-lapse microscopy in 35-mm glass-bottom culture dishes (GWST-3522, WillCo) in SC media at 30ºC essentially as described (Ferrezuelo *et al*, 2012), using a fully-motorized Leica AF7000 microscope with a 63X/1.3NA oil-immersion objective. Time-lapse images were analyzed with the aid of BudJ, an ImageJ (Wayne Rasband, NIH) plugin to obtain cell dimensions and fluorescence data as described (Ferrezuelo *et al*, 2012). Intracellular foci were detected with BudJ as pixels with a fluorescence value above a certain threshold relative to the median cell fluorescence that produced a contiguous area with a minimum size (both set by the user). In a typical set up, pixels were selected if at least 30% brighter than the cell median, with a minimal size of 0.2 μm. Photobleaching during acquisition was negligible (less than 0.1% per time point) and autofluorescence was always subtracted. Images z-stacks were obtained in a Zeiss 780 confocal microscope, and processed with ImageJ to produce average projections.

### Mutual-inhibition mathematical model

A wiring diagram and a set of differential equations were produced with the aid of COPASI (Hoops *et al*, 2006) to describe the mutual inhibition framework (Fig EV5A). First, we considered SG factors as either free components (*SG*_*Comp*_) or in a condensed state (*SG*), the stress signal (*Stress*) acting as a positive modulator in their condensation. Regarding SG dissolution, we assumed two independent mechanisms, a basal process that reverses condensation in the absence of stress, and a Cdk-dependent dissolution reaction, in which active Cdk (*Cdk*_*Cyc*_) acts as an enzyme with specific catalytic (*k*_*c*_) and Michaelis-Menten (*K*_*M*_) constants. In turn, active Cdk (*Cdk*_*Cyc*_) is inactivated (*Cdk*) as a function of condensed SG factors (*SG*) to simulate translation inhibition of G1 cyclin mRNAs by SGs. Finally, we assumed stress-independent reactivation kinetics of the Cdk. With the exception of Cdk-dependent SG dissolution, all processes were defined by mass-action laws with explicit rate constants. This simplicity allowed us to estimate all kinetic parameters from experimental data using COPASI in deterministic mode. First, the *K*_*M*_ for Cdc28 was previously determined *in vitro* (Bouchoux & Uhlmann, 2011), and the maximal substrate concentration in SGs (*SG*_*max*_) was estimated as described in Fig EV5B, allowing us to obtain a relative *K*_*Mr*_ =*K*_*M*_/*SG*_*max*_. We obtained the basal SG-dissolution rate constant (*k*_*bd*_) by fitting the model to SG dissolution in the *cdc28-13* mutant in the absence of stress (Fig EV5C), where all other variables have no effect. We then used SG formation and dissolution data from wild-type cells to estimate the remaining rate constants (Fig EV5D).

### Miscellaneous

Determination of mRNA levels by RT-qPCR and immunofluorescence analysis in yeast (Wang *et al*, 2004) and HeLa (Ruiz-Miro *et al*, 2011) cells were performed as described. Used antibodies are listed in Table EV3.

### Statistical analysis

Sample size is always indicated in the Fig legend. For single-cell or single-focus data, median and quartile (Q) values are shown. Pairwise comparisons were performed with a Mann-Whitney U test; and the resulting p-values are shown in the corresponding Fig panels. Time-lapse data from single cells during SG formation or dissolution are represented as the mean value of the population along time, while the shaded area represents the 95% confidence limits of the mean. Protein and mRNA levels were determined in triplicate samples and mean + SEM values are shown. Venn diagrams were generated with InteractiVenn (Heberle *et al*, 2015).

### Data and software availability

The model was deposited in the BioModels (Chelliah *et al*, 2015) database as MODEL2003140002 in SBML format together with a COPASI (Hoops *et al*, 2006) file to reproduce simulations with the parameter set shown in Fig EV5. BudJ (Ferrezuelo *et al*, 2012) can be obtained as an ImageJ (Wayne Rasband, NIH) plugin from ibmb.csic.es/groups/spatial-control-of-cell-cycle-entry.

---

**Figure EV1.**
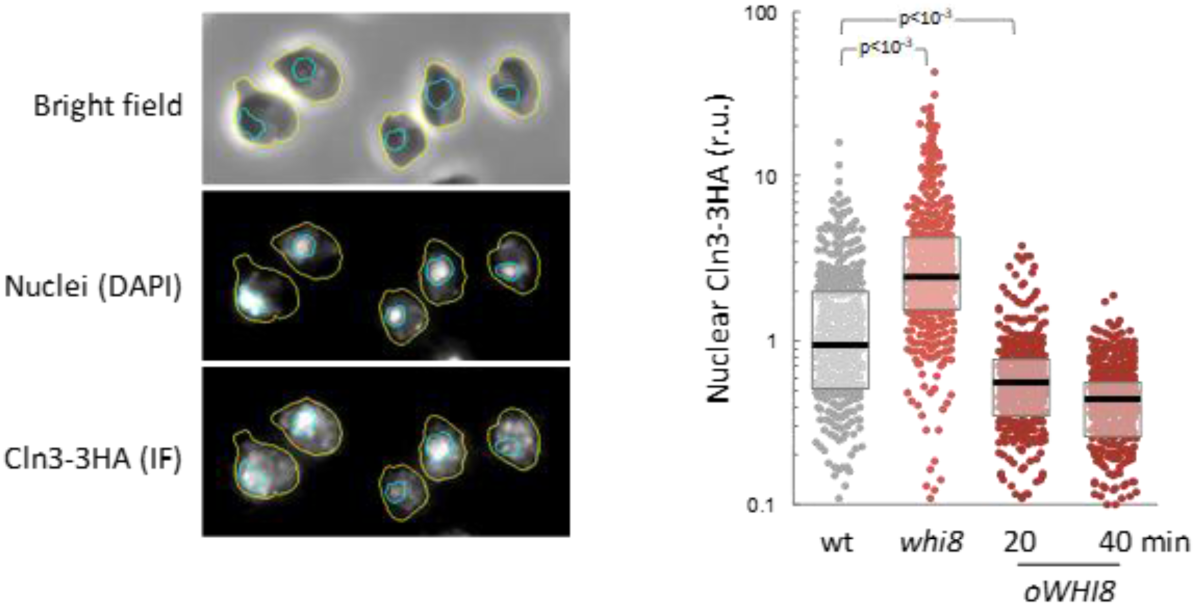
Whi8 counteracts nuclear accumulation of Cln3. Cells with the indicated genotypes were analyzed to determine Cln3-3HA nuclear levels by immunofluorescence (left, Vergés *et al*, 2007). Individual data (n>400) and median ± Q values are plotted (right). Shown p-values were obtained using a Mann-Whitney U test.

**Figure EV2.**
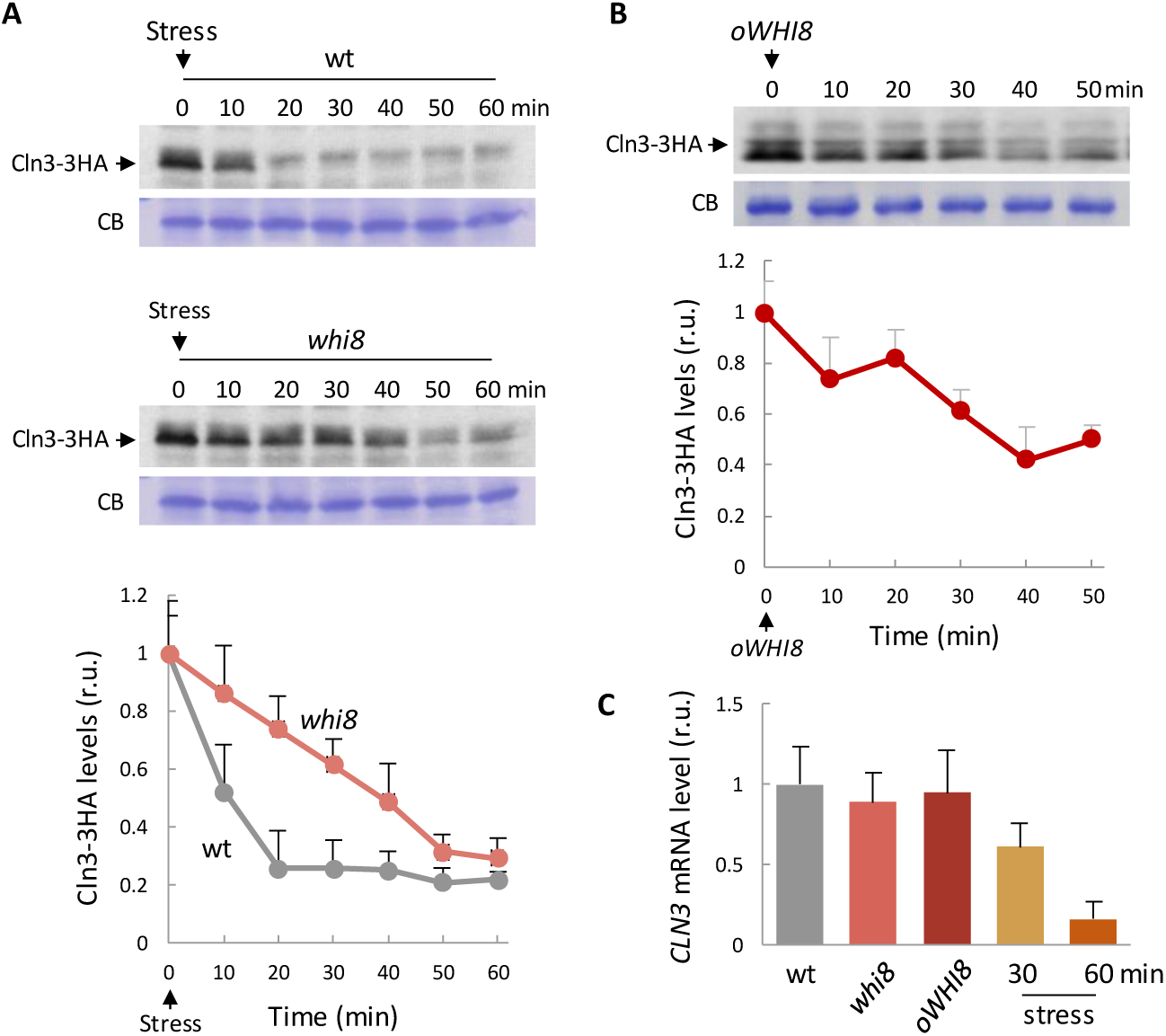
Whi8 is required to inhibit *CLN3* mRNA translation under stress conditions. A Immunoblot analysis of Cln3-3HA after transferring wild-type (top) or *whi8* (bottom) cells to 42ºC in the absence of glucose. Total cell extracts were stained with Coomassie blue and a prominent band is shown as loading control. Cln3-3HA levels were quantified and mean + SEM values (n=3) are plotted. B Immunoblot analysis of Cln3-3HA after 1 µM β-estradiol addition to induce a *GAL1p-WHI8* fusion in cells expressing the Gal4-hER-VP16 transactivator. A prominent band in the total cell extract is shown as loading control. Cln3-3HA levels were quantified and mean + SEM values (n=3) are plotted. C *CLN3* mRNA levels were determined in cells of the indicated genotypes under normal conditions, or 30-60 min after transfer to 42ºC in the absence of glucose. Mean + SEM values (n=3) are plotted.

**Figure EV3.**
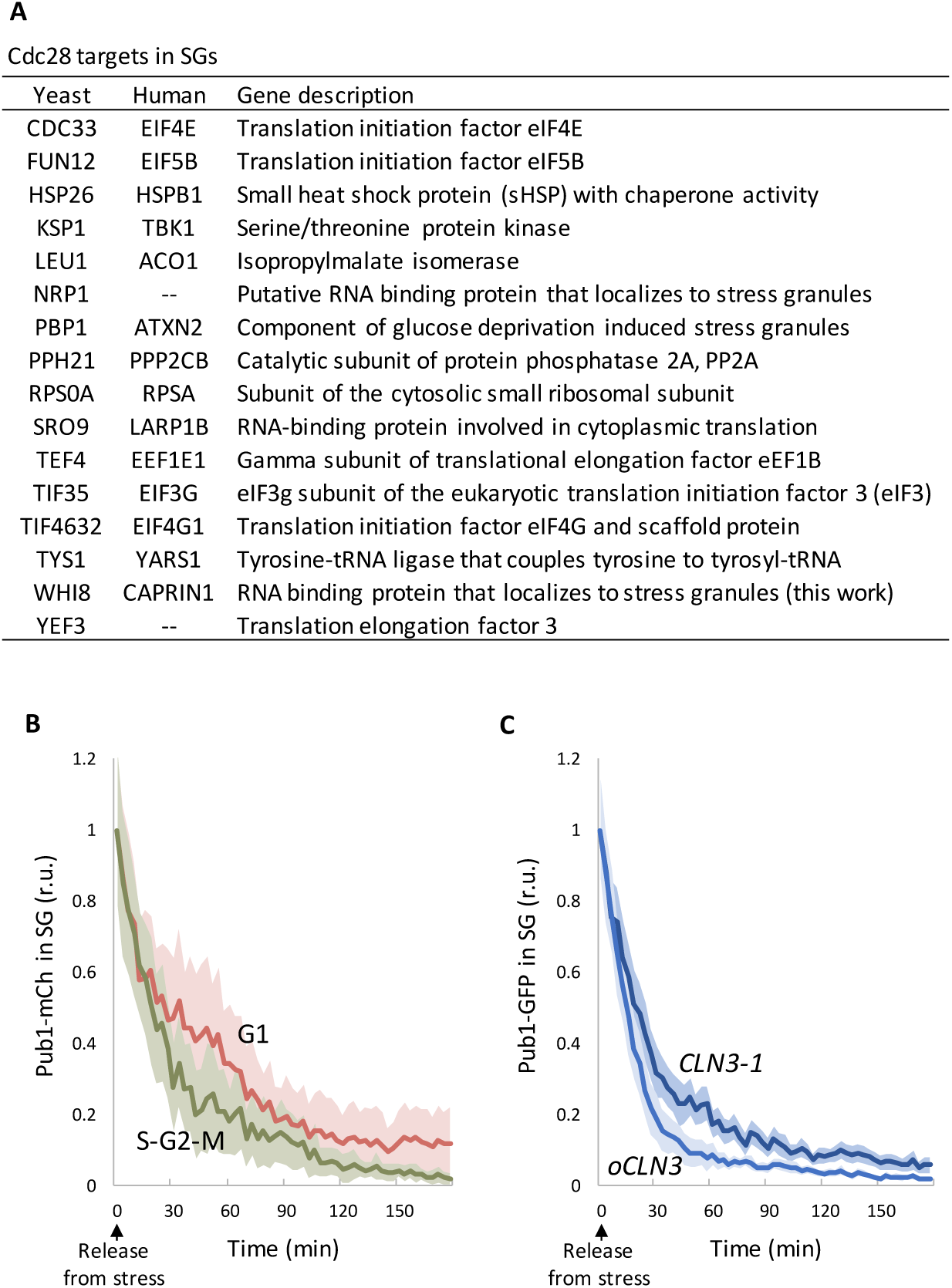
Cdc28 modulates SG dynamics. A Proteins in SGs with experimentally validated phosphosites complying with a Cdk consensus: (S/T)P with at least one basic amino acid from −2 to +3. B Wild-type cells expressing Pub1-mCherry were stressed for 30 min at 42ºC in the absence of glucose and, once released at 30ºC in the presence of glucose, Pub1-mCherry levels in foci were measured at different time points in unbudded (G1) or budded (S-G2-M) cells. Mean values (n>30) and confidence limits for the mean (α=0.05) are plotted. C Cells expressing Pub1-mCherry and a hyperstable allele (*CLN3-1*) or overexpressing wild-type Cln3 (*oCLN3*) from the *tetO_2_* promoter were stressed for 30 min at 42ºC in the absence of glucose and, once released at 30ºC in the presence of glucose, Pub1-mCherry levels in foci were measured at different time points. Mean values (n>30) and confidence limits for the mean (α=0.05) are plotted.

**Figure EV4.**
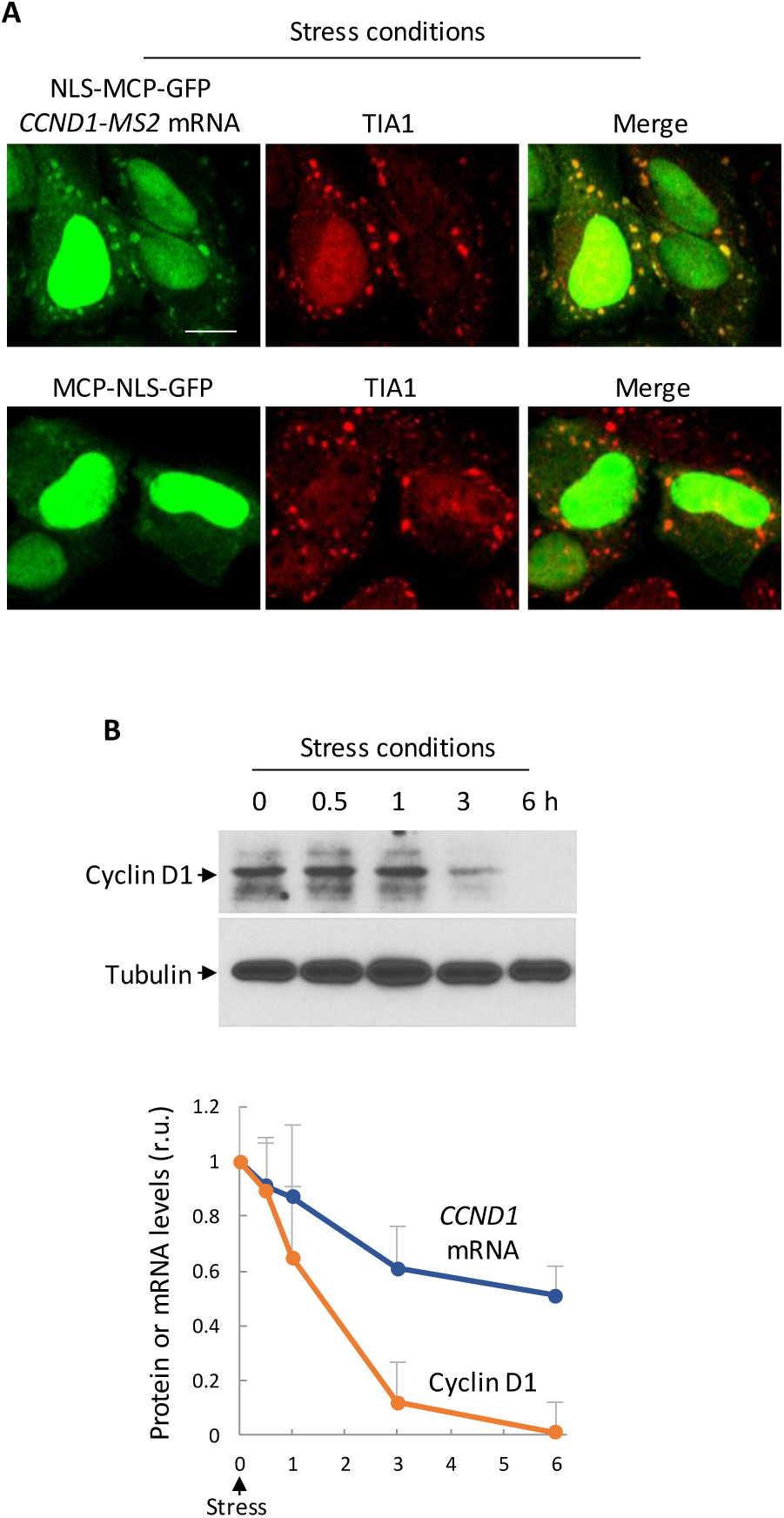
The *CCND1* mRNA is recruited to SGs and cyclin D1 levels are downregulated by stress. A HeLa cells expressing NLS-MCP-GFP and either a *CCND1-MS2* mRNA or none (as control) were subject to 0.5 mM NaAsO_2_ for 30 min, and analyzed by immunofluorescence with a αTIA1 antibody. Bar, 5 µm. B Immunoblot analysis of cyclin D1 in HeLa cells at the indicated times after addition of 0.5 mM NaAsO_2_. Tubulin is shown as loading control. Cyclin D1 protein and *CCND1* mRNA levels were quantified and mean + SEM values (n=3) are plotted.

**Figure EV5.**
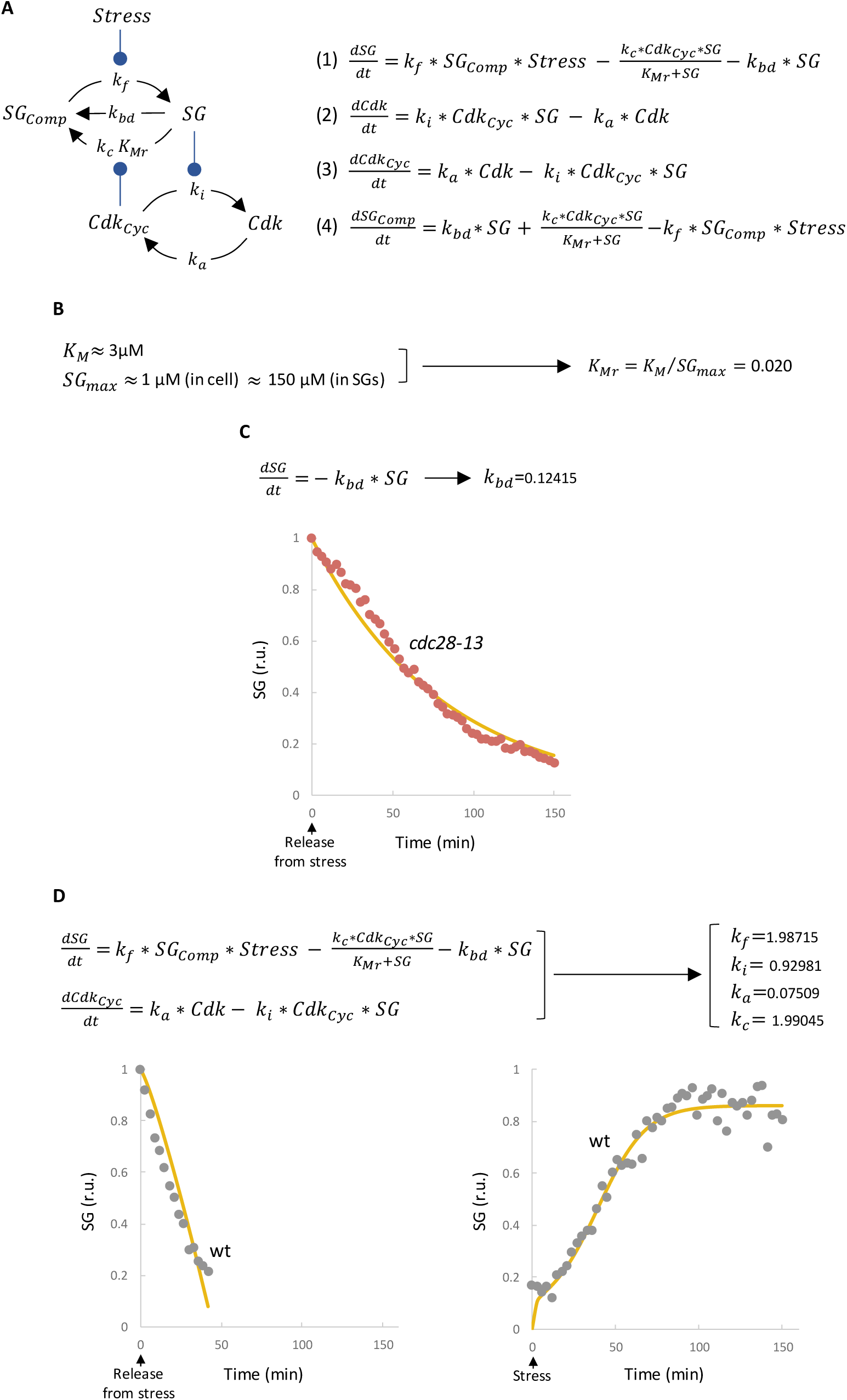
Equations and parameter fitting in the mutual-inhibition model. A Wire diagram of the mutual inhibition model opposing SG formation and Cdk activity. Variables and parameters used in the model are indicated. This model has four state variables: *SG*_*Comp*_, SG free components; *SG*, SG condensed factors; *Cdk*_*Cyc*_, active Cdk-cyclin complexes; and *Cdk*, inactive Cdk molecules. With the exception of SG dissolution by Cdk, all reactions are driven by simple mass-action laws with explicit parameters. Stress acts on SG formation by modulating condensation of SG components with *k*_*f*_, the formation rate constant. SG dissolution, in turn, takes place through (1) a default basal process with rate constant *k*_*bd*_, and (2) a Cdk-mediated enzymatic mechanism with *k*_*c*_ (catalytic constant) and *K*_*Mr*_ (relative Michaelis-Menten constant) parameters. Cdk is activated at a constant rate (*k*_*a*_) and, as cyclin mRNA becomes translationally inhibited, is downregulated by SGs with an inactivation constant *k*_*i*_. The set of non-linear differential equations used to simulate the model is also shown. B Cdc28 has a mean *K*_*M*_≈3 μM (Bouchoux and Uhlmann, 2016), and putative Cdc28 targets in SGs (Fig EV3A) display an average concentration close to 1 μM (Ho *et al*, 2018). We carefully analyzed Whi8-GFP and Pub1-GFP levels (as in Fig 4A) and estimated that these proteins increase their concentration by ca. 150-fold in SGs under stress conditions. Thus, *K*_*Mr*_ = *K*_*M*_/*SG*_*max*_ ≈ 0.02. C The basal SG-dissolution rate constant (*k*_*bd*_) was obtained by fitting the model to SG dissolution in the *cdc28-13* mutant in the absence of stress, where all other variables have no effect. Equations used and the resulting fitted curve (yellow line) are shown. D The remaining parameters of the model were obtained by fitting the model to SG formation and dissolution experimental data (gray points) from wild-type cells. Equations used and the resulting fitted curves (yellow lines) are shown.

**Table EV1.**
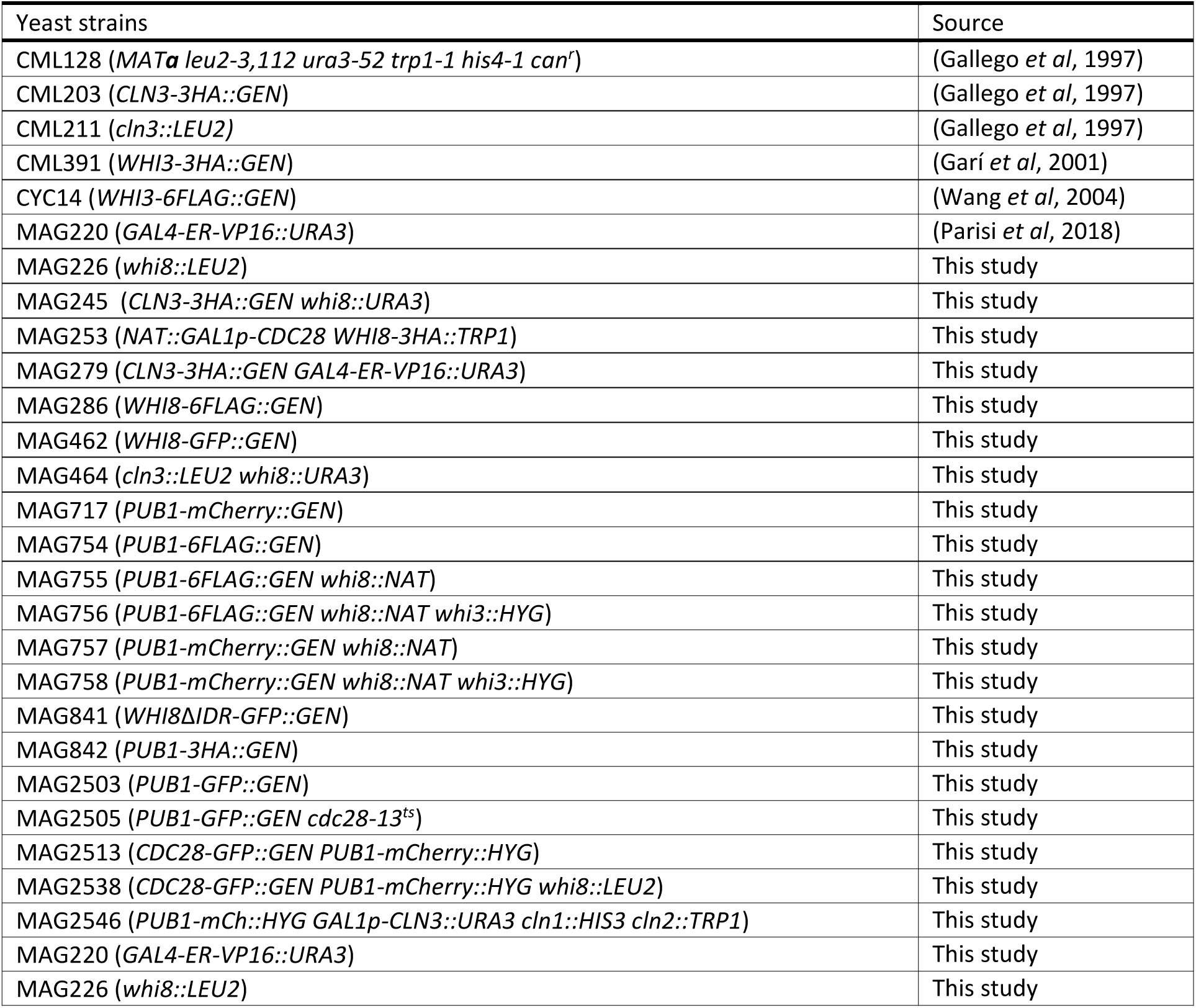
Yeast strains.

**Table EV2.** Plasmids.

| Plasmids | Source |
| --- | --- |
| pCM192 (*ARS-CEN URA3 CLN3-1-3HA*) | (Gallego *et al*, 1997) |
| pCM194 (*ARS-CEN URA3 CLN3-3HA*) | (Gallego *et al*, 1997) |
| pCM255 (*ARS-CEN URA3 tetO2-CLN3-3HA tTA*) | (Gallego *et al*, 1997) |
| pCYC22 (2μm *URA3 GAL1P-cln3<sup>nd</sup>-MS2*) | This study |
| pCYC2077 (*UBIp-3HA-CDK4*) | (Ruiz-Miro *et al*, 2011) |
| pG14-MS2-GFP (2μm *LEU2 GPDp-NLS-MCP-GFP*) | (Bertrand *et al*, 1998) |
| pMAG140 (*ARS-CEN LEU2 CDC28-3FLAG*) | (Yahya *et al*, 2014) |
| pMAG141 (*ARS-CEN LEU2 CDC28<sup>wee</sup>-3FLAG*) | (Yahya *et al*, 2014) |
| pMAG231 (*ARS-CEN TRP1 GAL1p-WHI8*) | This study |
| pMAG752 (2μm *LEU2 GPDp-NLS-MCP-mCherry*) | This study |
| pMAG786 (*CMVp-CCND1-MS2*) | This study |
| pMAG788 (*CMVp-NLS-MCP-GFP*) | This study |
| pMAG1307 (*ARS-CEN LEU2 WHI8-6FLAG*) | This study |
| pMAG2491 (*CMVp-mCherry-G3BP1*) | This study |

**Table EV3.** Antibodies

| Antibodies | Source | Identifier |
| --- | --- | --- |
| αCaprin1 | Proteintech | 15112-1-AP |
| αCcnD1 | Upstate | AB_1977143 |
| αDpm1, clone 5C5A7 (mouse) | Invitrogen | A6429 |
| αFLAG, clone M2 (mouse) | Sigma | MAB2200 |
| αGFP, clones 7.1 and 13.1 (mouse) | Roche | 11814460001 |
| αHA, clone 12CA5 (mouse) | Roche | 11583816001 |
| αHA, clone 3F10 (rat) | Roche | 11867423001 |
| αHxk1 | US Biological | H2035-03 |
| αTIA1 | Santa Cruz Biotechnology | sc-1751 |

